# Generation of *C9orf72* repeat knock-in iPSC lines for modelling ALS and FTD

**DOI:** 10.1101/2025.02.10.637041

**Authors:** Rachel Coneys, Alexander J. Cammack, Remya R. Nair, David Thompson, Jonas Mechtersheimer, Mireia Carcolé, Yashica Gupta, Gabriel E. Rech, Michael Flower, Niamh O’Brien, Marc-David Ruepp, Sarah Mizielinska, Fiona Ducotterd, Sarah J. Tabrizi, Elizabeth M.C. Fisher, Thomas J. Cunningham, Michael Ward, William C. Skarnes, Adrian M. Isaacs

## Abstract

Induced pluripotent stem cell (iPSC) models are powerful tools for neurodegenerative disease modelling, as they allow mechanistic studies in a human genetic environment and they can be differentiated into a range of neuronal and non-neuronal cells. However, these models come with inherent challenges due to line-to-line and clonal variability. To combat this issue, the iPSC Neurodegenerative Disease Initiative (iNDI) has generated an iPSC repository using a single clonal reference line, KOLF2.1J, into which disease-causing mutations and revertants are introduced via gene editing. Here we describe the generation and validation of lines carrying the most common causative mutation for amyotrophic lateral sclerosis (ALS) and frontotemporal dementia (FTD), a repeat expansion in the *C9orf72* gene, for the iNDI collection of neurodegenerative iPSC models. We demonstrate that these *C9orf72* knock-in lines differentiate efficiently into neurons and display characteristic *C9orf72*-associated pathologies, including reduced C9orf72 levels and the presence of dipeptide repeat proteins (DPRs) and RNA foci, which increase in abundance over time in culture. These pathologies are not present in revertant cells lacking the repeat expansion. These repeat expansion and revertant cell lines are now available to academic and for-profit institutions through the JAX iPS cell repository and will help to facilitate and standardise iPSC-based ALS/FTD research.

## Introduction

A GGGGCC hexanucleotide repeat expansion in the first intron of *C9orf72* is the most common genetic cause of amyotrophic lateral sclerosis (ALS) and frontotemporal dementia (FTD) (C9 ALS/FTD) [1,2]. Three non-mutually exclusive mechanisms have been proposed to be responsible for *C9orf72*-associated toxicity [3]: haploinsufficiency of the C9orf72 protein; toxic repeat-containing RNAs; and toxic dipeptide repeat proteins (DPRs) generated by repeat-associated non-ATG (RAN) translation. Induced pluripotent stem cells (iPSCs) derived from ALS/FTD patients and their isogenic controls have been fundamental to our understanding of disease mechanisms [4] and large-scale community efforts have developed valuable resources of patient-derived lines [5,6]. However, one challenge with iPSCs derived from different donors is the inherent variability between lines and therefore use of standardised iPSC models would complement patient iPSCs. To standardise the iPSC lines used across the whole neurodegenerative disease community, the iPSC Neurodegenerative Disease Initiative (iNDI) has generated a collection of neurodegeneration-relevant iPSC models, established in a single wild-type reference cell line, KOLF2.1J [7,8]. Using genome editing, disease-causing mutations have been introduced at their endogenous loci, allowing for cross-analysis of different mutations in the same genetic background. Revertant lines, in which the mutations are corrected back to the non-disease allele, were also generated, creating a crucial control allowing any observed changes to be more readily attributed to the disease mutation, rather than the editing process or clonal selection. Therefore, the use of a ‘trio’ of lines (parental, mutated, revertant) for each familial mutation simplifies establishing whether phenotypes are driven by the disease-causing mutation.

The *C9orf72* repeat expansion represents a unique challenge for this approach due to its length and GC content, which make it hard to manipulate through traditional cloning methods, as well as its well-documented somatic instability [9]. Here, we detail the generation and characterisation of *C9orf72* repeat expansion knock-in lines in the KOLF2.1J background to be included within the iNDI catalogue of neurodegenerative models. Targeting constructs harbouring large pathogenic repeat expansions were engineered via a bespoke cloning pipeline originally developed for generation of animal models [10]. These were subsequently reengineered with human homology arms enabling CRISPR-Cas9 assisted homologous recombination at the endogenous human *C9orf72* locus. Two distinct heterozygous *C9orf72* repeat knock-in lines were generated with seamless expanded G_4_C_2_ repeats, and then edited again to remove the repeat expansion to generate two companion revertant lines. We show that these novel *C9orf72* knock-in lines differentiate efficiently into neurons, display reduced C9orf72 levels, and harbour DPRs and RNA foci which increase in abundance as the cells age. These lines are now freely available as a resource to the ALS/FTD research community to facilitate future studies.

## Results

### Generation of *C9orf72* repeat knock-in iPSC lines

The strategy for seamless knock-in of G_4_C_2_ hexanucleotide repeats into one copy of the *C9orf72* gene is shown in Figure 1. We first engineered a 350 G_4_C_2_ repeat targeting vector (Supplementary Figure 1) in the pJazz backbone [11] where the repeats are flanked by 1 kb (5’) and 2.9 kb (3’) homology arms consisting of endogenous human sequence (Figure 1A), which we subsequently used for Cas9-mediated targeting in KOLF2.1J iPSCs. Taking advantage of error-prone non-homologous end joining (NHEJ) repair at Cas9-induced double-stranded breaks, we began by creating a heterozygous indel (8 bp deletion) in intron 1 using Cas9 ribonucleoprotein (RNP), thereby creating a novel CRISPR site in one of the two alleles of *C9orf72* (Figure 1B). This was followed by co-nucleofection of the linear double-stranded 350-repeat pJazz donor vector and Cas9 RNP targeted to this novel sequence (Figure 1C), performed under optimized conditions that promote high-efficiency homology-directed repair [12]. Ninety-six single-cell derived clones were initially screened by repeat-primed PCR (rpPCR) to identify clones carrying at least 35 hexanucleotide repeats. Southern blotting identified two clones, D08 and F05, that contained ~200 hexanucleotide repeats (Supplementary Figure 2A). These clones were expanded for further characterization and subsequently used for revertant experiments. To revert the repeat expansion, D08 and F05 cells were first nucleofected with Cas9 RNP complexed with two guide RNAs designed to remove the repeat and some flanking sequence (Figure 1D). These cell pools were then nucleofected with a plasmid donor containing wild-type C9 sequence and a guide RNA designed to match the predicted deletion junction formed by perfect NHEJ (Figure 1E, F). Following single cell cloning, PCR amplification and Sanger sequencing across the target region identified revertant clones (D08 C03 and F05 B07) with two intact copies of wild-type C9 sequence, inferred by the presence of a nearby KOLF2.1J-specific SNP.

**Figure 1.**
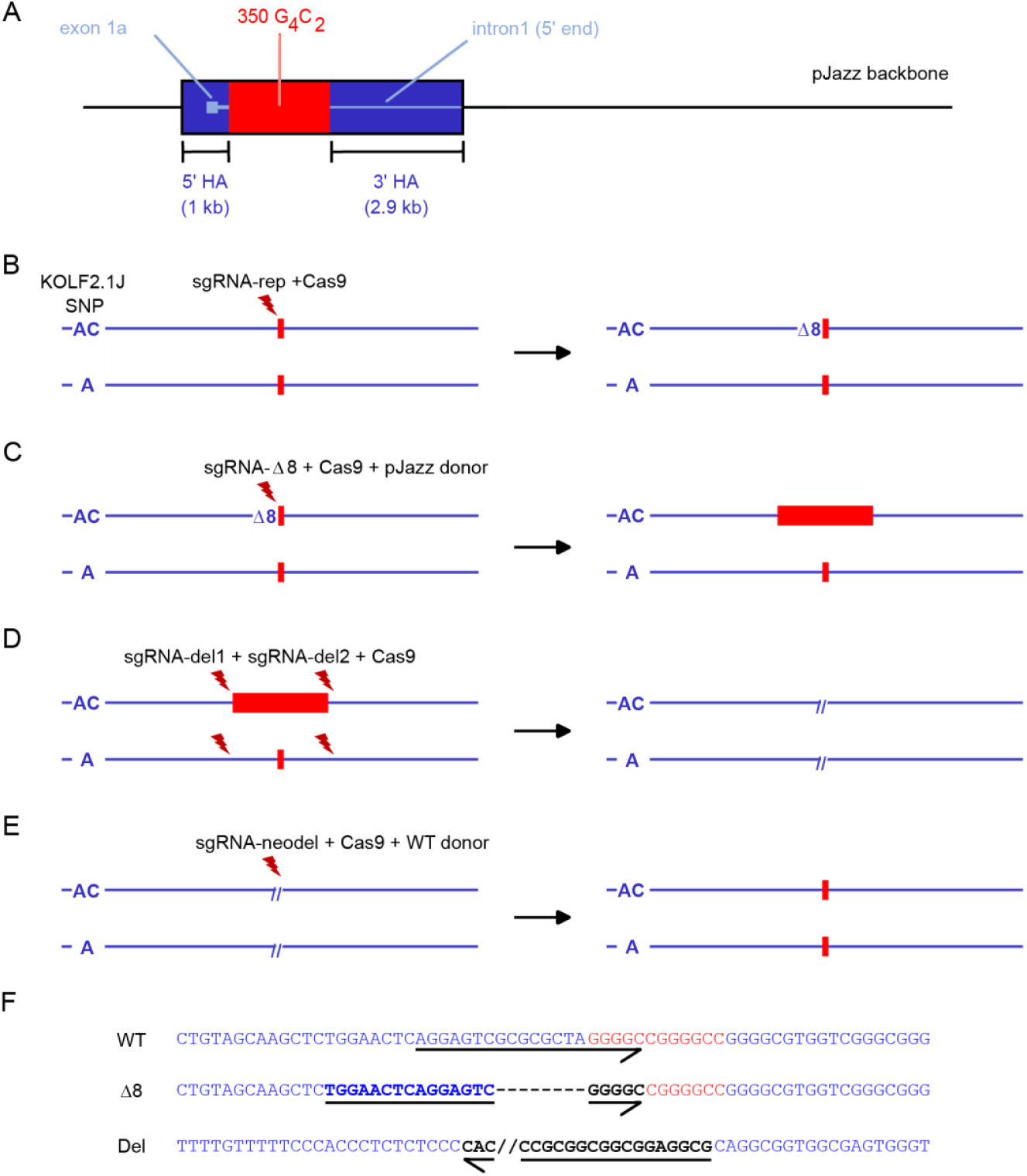
Strategy for engineering of *C9orf72* hexanucleotide G_4_C_2_ repeat and revertant alleles. (**A**) pJazz donor plasmid containing 350 copies of the hexanucleotide repeat flanked by *C9orf72* homology arms (HA). (**B-E**) Stepwise generation of allele-specific repeat expansion and reversion cell lines. (B) Introduction of an 8 base pair deletion (Δ8) in one copy of the *C9orf72* gene by NHEJ repair Cas9 RNP and sgRNA-rep. (C) Insertion of G_4_C_2_ repeats by homology-directed repair with pJazz donor. (D) Removal of repeat region and flanking sequence in both copies of *C9orf72*. (E) Repair of deletion by homology-directed repair with WT donor plasmid containing wild-type *C9orf72* sequence. (F) Sequence of wild-type (WT), allele-specific indel (Δ8), and biallelic deletion (Del) alleles. Arrows underline the sequence of sgRNA-rep, sgΔ8, and sgRNA-neodel guide RNAs.

### C9 knock-in lines harbour the hexanucleotide repeat expansion

After expansion and archiving of the two isogenic pairs of knock-in and revertant C9 clones, rpPCR confirmed the presence of the repeat expansion in the F05 and D08 knock-in lines and its absence in the KOLF2.1J parental control and the revertant lines (Figure 2A). We then performed Southern blotting of genomic DNA in all five lines. F05 and D08 had approximately 200 repeats and these were absent in the revertant and parental KOLF2.1J lines (Supplementary Figure 2B). For accurate repeat-sizing and to confirm correct targeting, we performed Cas9-targeted long-read Oxford Nanopore Technology (ONT) sequencing of the entire *C9orf72* gene. Clones D08 and F05 had modal repeat sizes of 186 and 208 G_4_C_2_ repeats respectively (Figure 2B), with no alterations in the flanking sequence (Figure 2C and Supplementary Figure 3) Furthermore, G-band karyotyping and SNP array analysis [13] confirmed the knock-in and revertant clones retained a normal 46;XY karyotype and did not contain any additional copy number changes other than those found in the KOLF2.1J reference cell line (Supplemental Data File 1).

**Figure 2:**
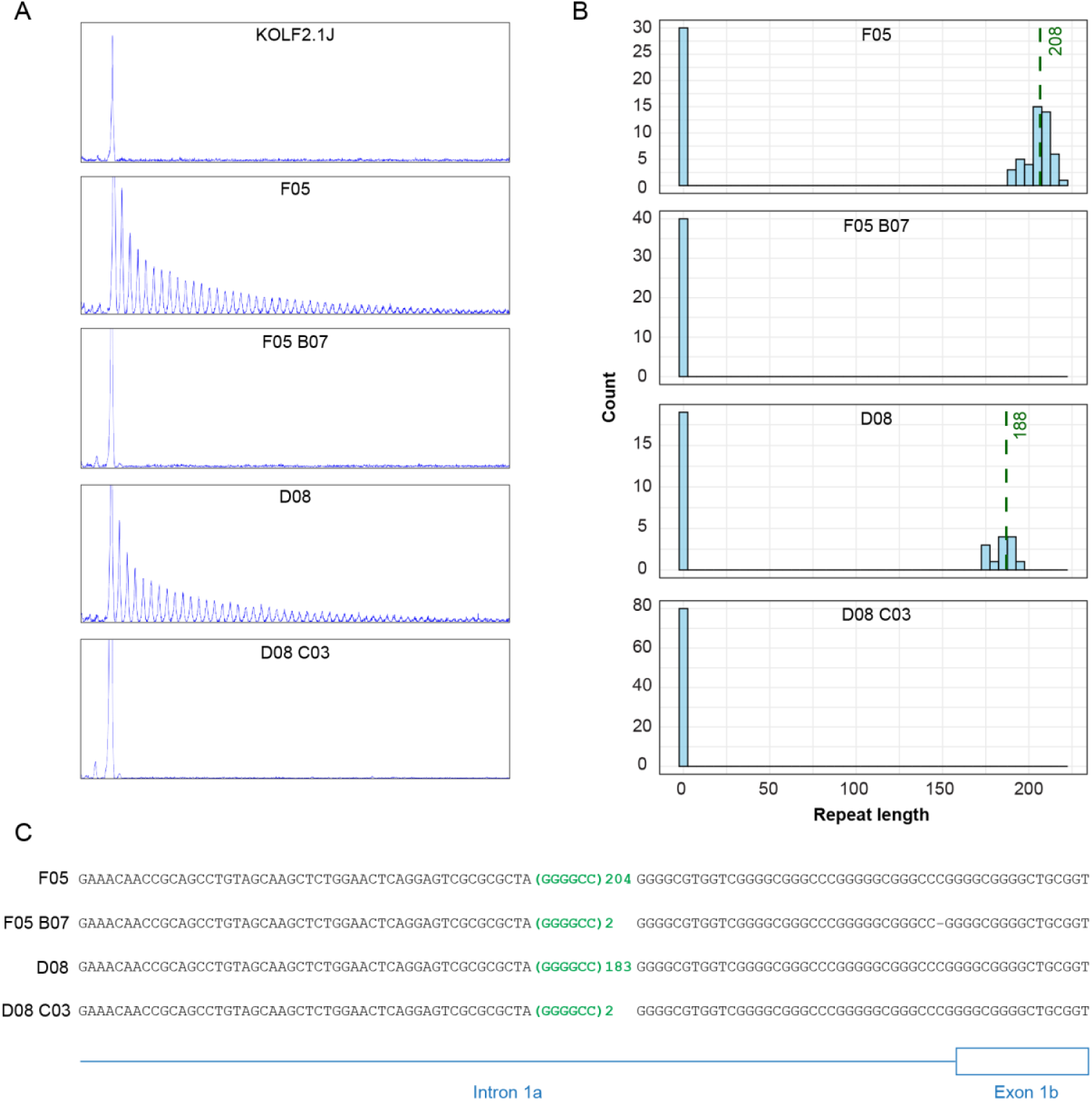
C9 knock-in lines contain expanded G_4_C_2_ repeats. (**A**) rpPCR of KOLF2.1J parental control, C9-knockin lines (F05 and D08), and revertant lines (F05-B07 and D08-C03). (**B**) Repeat length frequency distributions from Cas9-targeted ONT long-read sequencing. The x-axis represents hexanucleotide repeat length, and the y-axis the read count, with a bin width of 5 repeats. Dashed green lines represent modal repeat length. (**C**) ONT long-read consensus sequence flanking the repeats.

### C9 knock-in lines show reduced C9orf72 expression

*C9orf72* contains 11 exons which are alternatively spliced into 3 different splice variants (Figure 3A). The repeat expansion lies within intron 1 of variants 1 and 3, and within the promoter region of variant 2, which is the most abundant of the three [14]. In patients, a 25-50 % reduction of variants 1, 2 and total *C9orf72* mRNA and protein is seen in the frontal cortex and cerebellum [1,15–21] with an increase in intron 1a-containing transcripts [20]. In the knock-in iPSC lines, RT-qPCR showed only trends towards reduction of variant 2 and increased intron 1a transcripts (Figure 3B-C); however a significant reduction by approximately 50% was observed in C9orf72 protein levels in the F05 and D08 knock-in iPSC lines using western blotting, and this was reversed back to wildtype levels in the revertants (Figure 3D). Total *C9orf72* mRNA levels remained unchanged, likely balanced by the simultaneous increase in intron 1a transcripts and decrease in variant 2 (Figure 3B-C).

**Figure 3:**
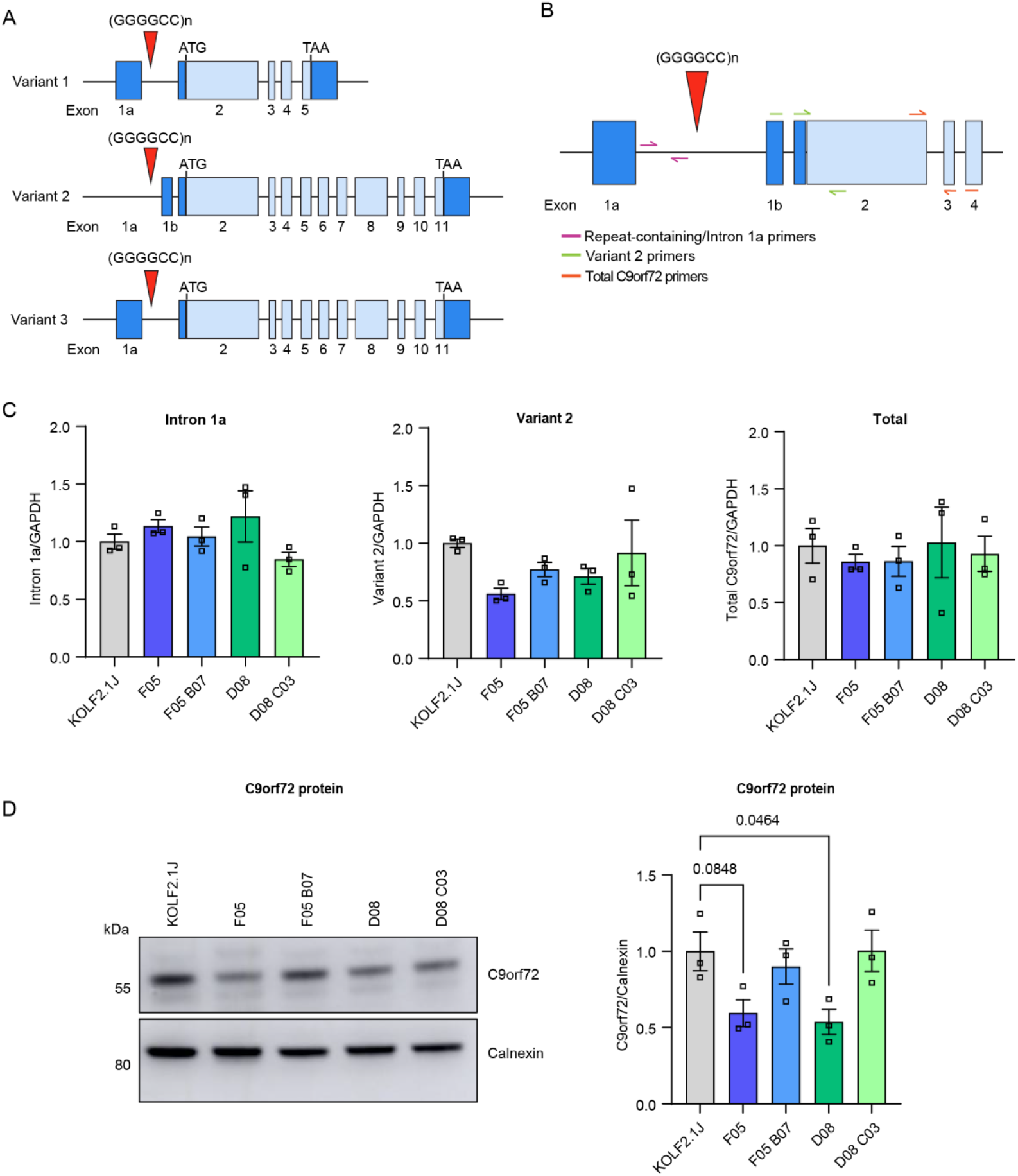
C9 knock-in iPSCs display reduced protein levels of C9orf72. (A) The three transcript variants of *C9orf72*. Untranslated exons are indicated by dark blue boxes and light blue boxes indicate coding exons. The repeat expansion is indicated by the red triangle, which is located between exons 1a and 2 of transcript variants 1 and 3, and within the promoter region of variant 2. (B) Schematic of RT-qPCR primer design. Primers specific for repeat-containing transcripts bind in intron 1a; primers specific for variant 2 bind across the exon1b – exon2 junction; primers detecting total *C9orf72* bind across exons 2 – exon 3–4. (C) RT-qPCR analysis of repeat-containing transcripts (intron 1a), variant 2, and total *C9orf72*, normalised to *GAPDH* in iPSCs, presented as fold-change over KOLF2.1J. Bars are mean ± SEM. N = 3 biological replicates per condition and n = 2 technical replicates per condition. (D) Representative immunoblot and quantification of C9orf72 and housekeeping control Calnexin in iPSCs. Bars are mean ± SEM. N = 3 biological replicates per condition and n = 2 technical replicates per condition.

### C9 knock-in lines differentiate into lower motor neurons with a similar efficiency as the KOLF2.1J parental line

To evaluate the presence of disease-specific pathology in neurons as well as iPSCs, we inserted an *NGN2* cassette into each line using piggyBac-mediated transposition to allow rapid differentiation into neurons (Figure 4A). We then differentiated the lines into inducible lower motor neurons (iLMNs) using *NGN2*-meditated differentiation coupled with small molecule patterning [22] (Figure 4A). All five lines differentiated efficiently, with 52-100 % of neurons in each line expressing motor neuron markers SMI-32, ISL1 and ChAT, and pan-neuronal marker MAP2 by day 10 (Figure 4B-C). RNA expression of the pluripotency marker OCT4 was dramatically reduced compared to iPSCs, while MAP2 and motor neuron markers HB9 and ChAT were increased in all lines to a similar degree (Figure 4D).

**Figure 4:**
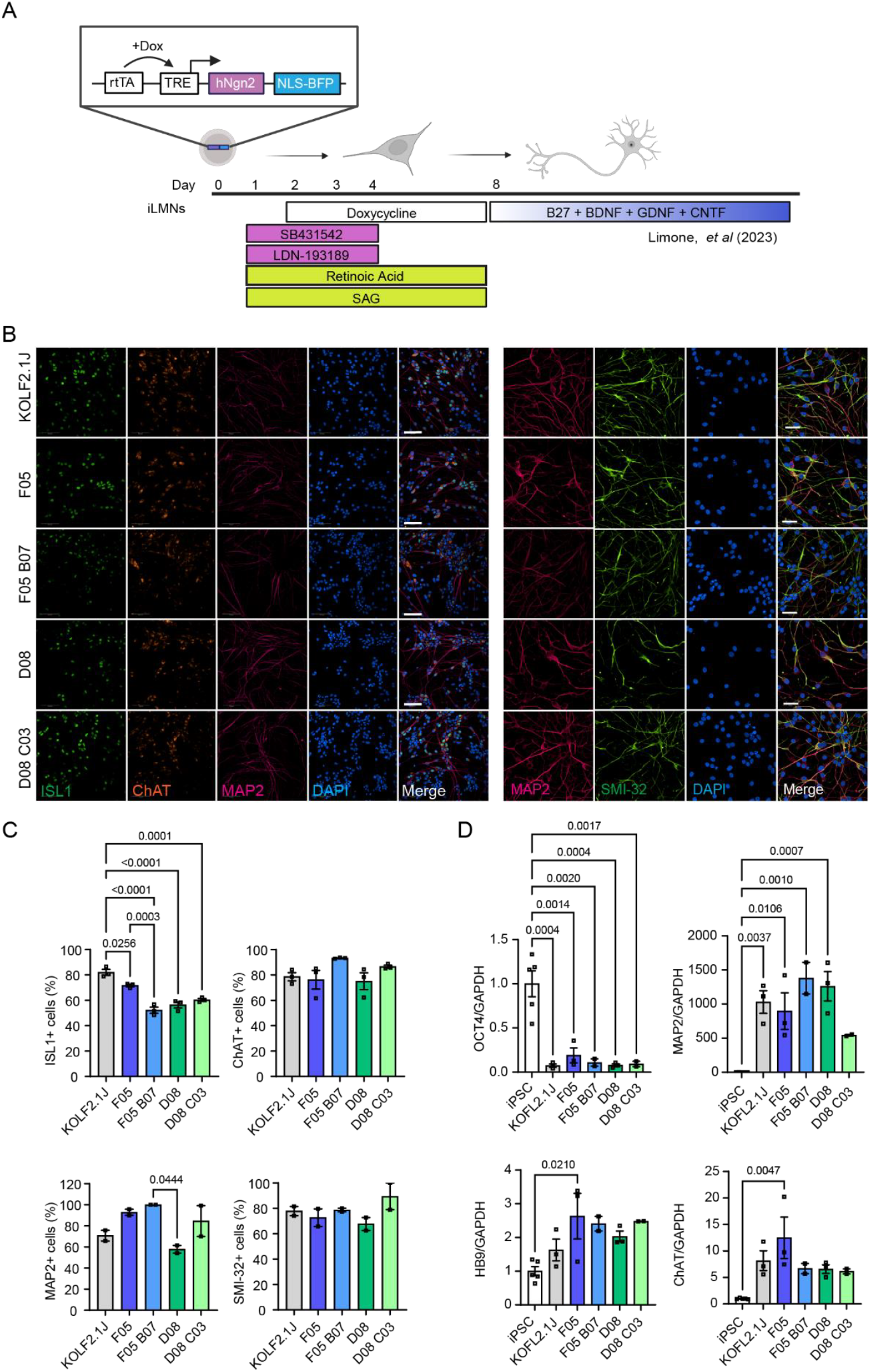
C9 knock-in lines differentiate efficiently into lower motor neurons. (A) Experimental process of the iLMN protocol [22]. Briefly, iPSCs stably expressing the *NGN2* construct are exposed to SB431542 and LDN-193189 on days 1-4, retinoic acid and SAG on days 1-8, and doxycycline on days 2-8, before media is changed to motor neuron maintenance media containing B27, BDNF, GDNF and CNTF. (B) Immunostaining of day 10 iLMNs; panel 1: MAP2 (pink), ChAT (orange), ISL1 (green), nuclei (blue), scale bar = 100 µm; panel 2: MAP2 (pink), SMI-32 (green) and nuclei (blue), scale bar = 50µm. (C) Quantification of ISL1, ChAT, MAP2 and SMI-32 positive cells in day 10 iLMNs. N = 2-3 biological replicates per condition and n = 3 technical replicates per condition. (D) *OCT4, MAP2, HB9* and *ChAT* expression normalised to *GAPDH* in day 10 iLMNs, presented as fold-change over iPSCs. Bars are mean ± SEM. N = 2-3 biological replicates per condition and n = 2 technical replicates per condition.

### Repeat RNA foci are detected in the C9 knock-in lines and increase with differentiation and maturity

The accumulation of repeat RNA into RNA foci is a well-described pathology associated with the *C9orf72* repeat expansion [23]. We assessed the presence of sense RNA foci in iPSCs and day 10 iLMNs. As predicted, knock-in of the repeat expansion led to substantial RNA foci production (Figure 5A), which increased from being present in ~37 % and 28 % of F05 and D08 iPSCs, to ~74 % and 78 % in day 10 iLMNs (Figure 5B). Correspondingly, the number of foci per nuclei also increased with neuronal differentiation, increasing from an average of ~0.3 foci per nuclei in iPSCs to ~2.1 and ~2.6 foci per nuclei in F05 and D08 day 10 iLMNs (Figure 5C).

**Figure 5:**
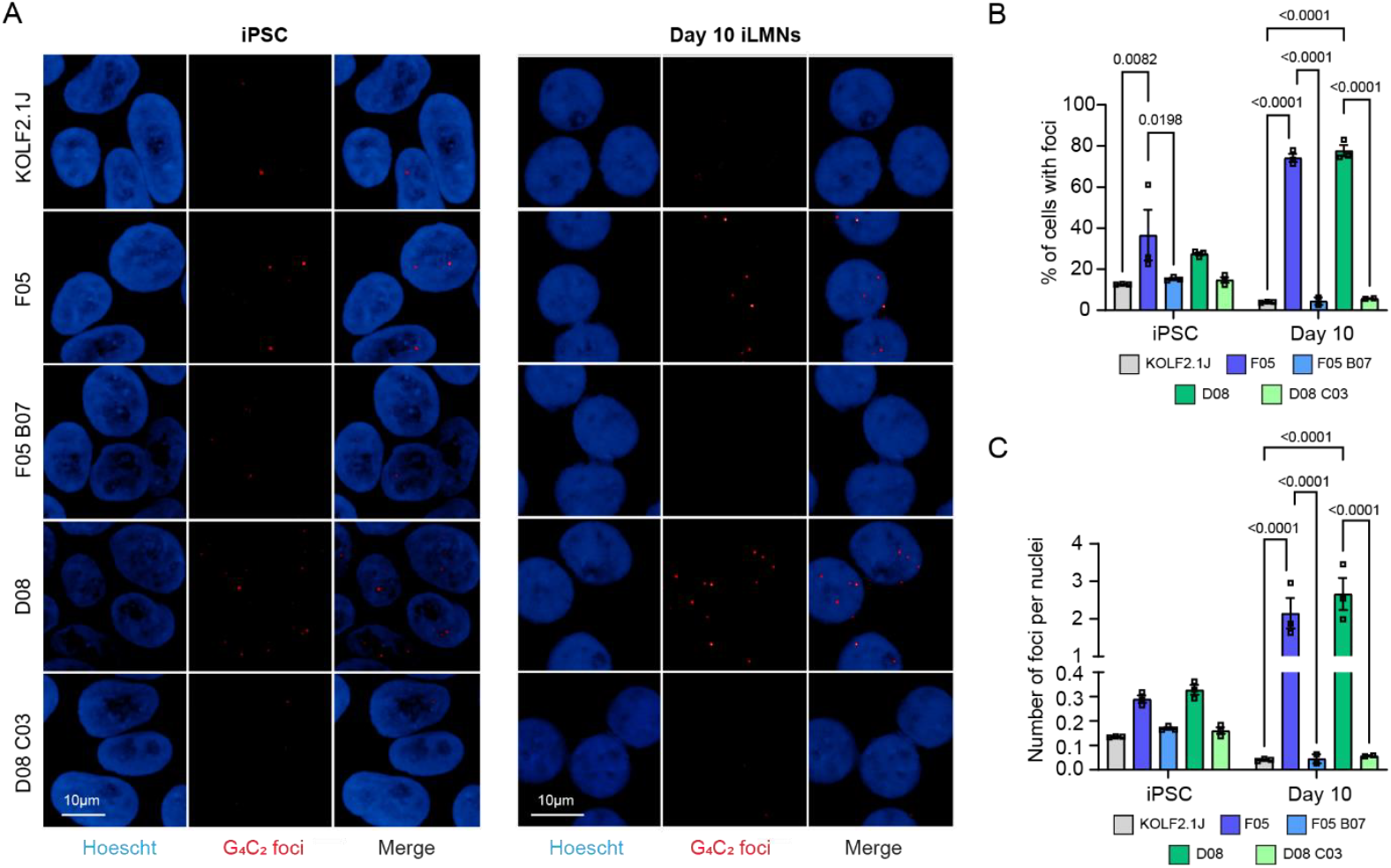
C9 knock-in lines display RNA foci which increase in abundance over time in culture. (A) RNA-FISH for sense G_4_C_2_ transcripts in iPSCs and day 10 iLMNs. Scale bar = 10 µm. (B) Percentage of cells with foci. Bars are mean ± SEM. N = 3 biological replicates per condition and n = 2-3 technical replicates per condition. (C) Number of foci per nuclei. Bars are mean ± SEM. N = 3 biological replicates per condition and n = 2-3 technical replicates per condition.

### PolyGA and polyGP are detected in the C9 knock-in lines and increase with differentiation and maturity

We next measured DPR levels in the C9 knock-in lines over time, from iPSCs to day 28 iLMNs. PolyGA was detected in iPSCs, day 10 and day 28 iLMNs in both C9 knock-in lines (Figure 6A), while polyGP was detected at all three timepoints in clone D08 and emerged by day 10 in clone F05 (Figure 6B). Since DPR burden was highest at day 28, we repeated polyGA and polyGP measurements at this timepoint with inclusion of the revertant lines. DPRs were again highly expressed in F05 and D08, but we observed only background signal in KOLF2.1J and the revertant lines, confirming DPRs are specific to the presence of the inserted repeat expansion (Figure 6C-D).

**Figure 6:**
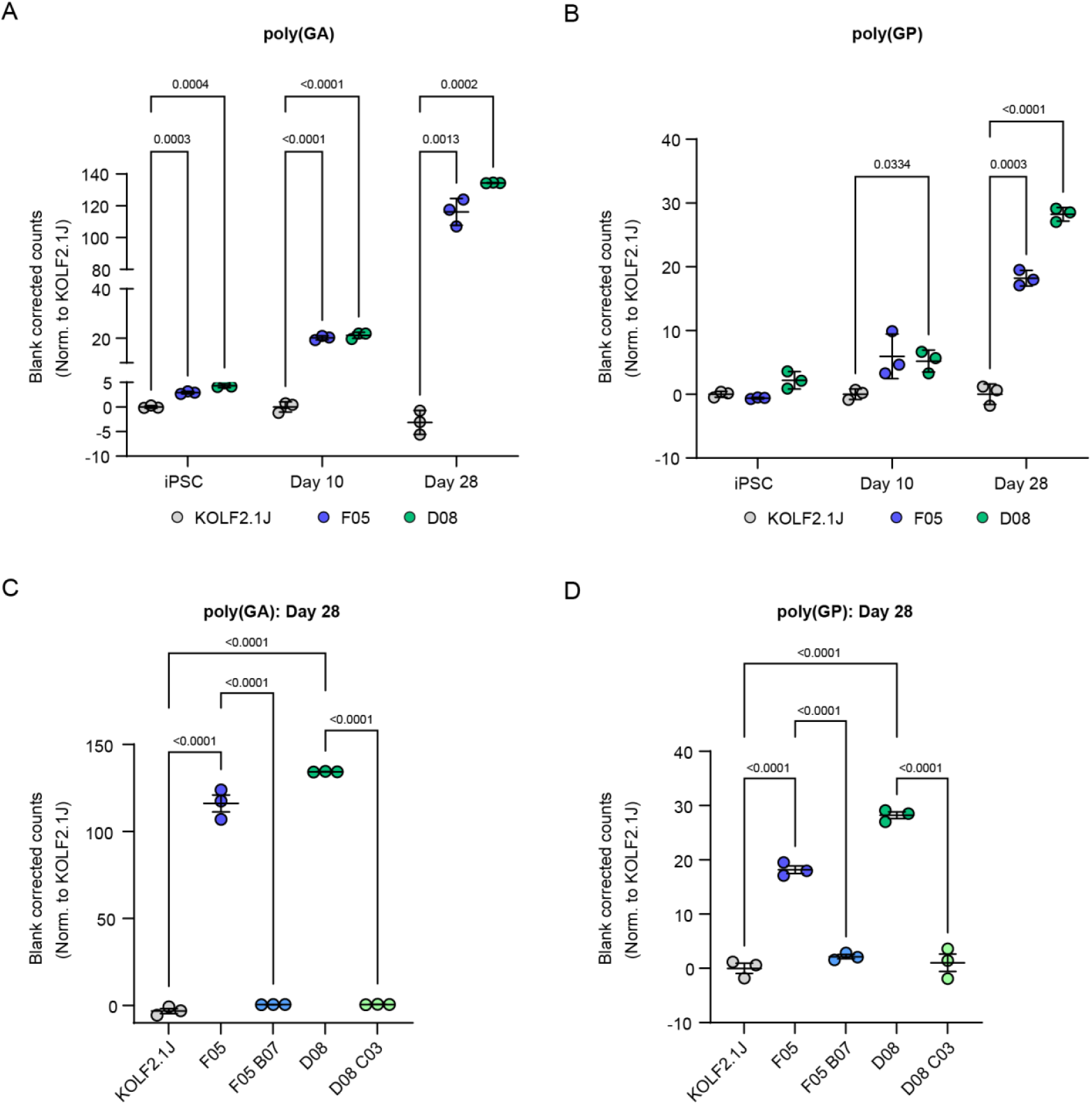
DPRs are detected in C9 knock-in lines and increase over time. (A-B) poly(GA) (A) and poly(GP) (B) blank corrected counts in KOLF2.1J, F05 and D08 iPSCs, day 10 and day 28 iLMNs, presented as fold change over KOLF2.1J at each timepoint. (C-D) poly(GA) (C) and poly(GP) (D) blank corrected counts in day 28 iLMNs compared to C9 revertant lines. Bars are mean ± SEM. N = 3 biological replicates per condition and n = 2 technical replicates per condition.

## Discussion

To provide a standardised model for the research community that helps address clonal variability and phenotypic heterogeneity within C9 ALS/FTD patient lines, we generated heterozygous C9 repeat knock-in lines, on the common reference background line, KOLF2.1J. We demonstrated that C9 knock-in lines differentiate efficiently into ALS-relevant iLMNs, contain the repeat expansion within the pathological range, and display the characteristic C9 pathologies of reduced C9orf72 expression, RNA foci and DPRs. We further show that RNA foci and DPR burden increases with time during neuronal differentiation. Critically, we successfully removed the repeat expansion to create revertant control lines and show these lines are absent of C9 pathologies. Thus, this set of lines is ready for use in the study of C9 ALS/FTD.

After extensive characterisation, KOLF2.1J was chosen as the iNDI reference line, largely based on its superior fast growth kinetics, amenability to CRISPR-Cas9 editing, and efficiency in neuronal differentiation [7]. The generation of these C9 knock-in lines adds to the rapidly growing collection of disease mutation lines on the KOLF2.1J background that are curated through the iNDI initiative. We now provide inclusion of the most common ALS/FTD mutation, which will allow comparisons across ALS/FTD mutations on a common background. However, we acknowledge there are limitations to assessing disease specific biology in a single genetic background from a healthy donor, and so we anticipate use of these standardised lines alongside C9-patient iPSCs, such as those generated by the Answer ALS consortium [5].

A crucial aspect of this model was the creation of revertant lines in which the repeat expansion is excised back out, restoring the C9 knock-in line back to the parental state. This will aid substantially in determining whether any observed phenotype is truly driven by the repeat expansion, as opposed to clonal variability which can hinder studies using patient lines and isogenic controls. In our revertant clones, DPRs and sense foci were no longer present following excision of the repeat expansion, confirming successful removal of the repeat expansion. Use of these revertants alongside the C9 knock-in lines provides an important and helpful new tool for the ALS/FTD research community. These lines are now ready for use in ALS/FTD research and are readily available to researchers through the iNDI initiative.

## Methods

### Donor plasmid construction

A targeting vector in the pJazz backbone [11] was engineered harbouring 350 G_4_C_2_ repeats flanked by 1 kb (5’) and 2.9 kb (3’) homology arms consisting of endogenous human sequence [10] as described in Supplementary Figure 1.

### iPSCs and CRISPR-Cas9 editing

KOLF2.1J cells [7] were cultured on Synthemax or Matrigel-coated plates in complete StemFlex media and nucleofected with Cas9 RNP as described previously [12].Plasmid DNA donors were ethanol precipitated and resuspended in P3 primary buffer prior to use. In experiments involving a donor template, nucleofected cells were plated in StemFlex media containing ROCK inhibitor (1X Revitacell; Invitrogen) and HDR enhancer v2 (1 μM; IDT) for 24 hours and incubated at 32°C (cold shock conditions) for 3 days.

### Cas9-targeted Nanopore sequencing and bioinformatic analysis

Two pairs of single-guide RNA (sgRNA’s) were designed up and down stream of the *C9orf72* locus avoiding any common SNPs (GTCTGTACTTAGTTTCCGCA, ATAGTAGTCAACTTAAGTAA, CGCCTAGTCCGTAATGATTA, CACCTAGAAGTACCTCAGGT). Prior to genomic DNA extraction (gDNA) tissue was pulverized on dry ice. All gDNA extraction was conducted using Monarch® HMW DNA Extraction Kit for Tissue (New England Biolabs). CRISPR-Cas9 targeted libraries were constructed as detailed in [24]. The resulting targeted library was sequenced for 48 hours on a GridION using a R9.4.1 flow cell (Oxford Nanopore Technologies). Basecalling was performed using GUPPY (v7.1.4). The resulting fastq files were trimmed using Porechop (v0.2.4) and quality controlled using NanoPlot (v1.29.1). A custom reference sequence was constructed based on the Ensembl *C9orf72* reference (ENSG00000147894), incorporating the two regions flanking the hexanucleotide repeat (available at https://github.com/mike-flower/othello.git). Reads were aligned to this custom reference using minimap2 with the ‘map-ont’ preset. Alignment SAM files were converted to sorted and indexed BAM files using samtools. Alignment quality was assessed in R by evaluating mapping quality, analysing the CIGAR strings, and calculating alignment length, mismatch, and indel counts. High-confidence alignments were selected based on mapping quality scores and consistent directionality. Antisense strands were reoriented to the sense strand to maintain consistency. Following alignment, custom R functions were used to detect hexanucleotide repeat motifs. Gaps between repeat units were allowed to account for sequence variants and sequencing artifacts. The repeat length was determined as the longest continuous stretch of repeat units, requiring at least a perfect repeat at both the start and end of the tract. Waterfall plots were generated in R using ggplot2 to visualise each read in relation to the *C9orf72* repeat region, including a 200-base window on either side of the repeat. The consensus sequence for each cluster was generated by performing multiple sequence alignment (MSA) using a custom R function that incorporates the msa package with the ClustalW algorithm. Pairwise alignment scores were calculated for each sequence against the consensus, and outliers were identified based on standard deviation from the mean alignment score. The bioinformatics pipeline is available at https://github.com/mike-flower/othello.git.

### iPSC culture for neuronal differentiations

iPSCs were maintained on Geltrex-coated plates in StemFlex media supplemented with StemFlex supplement, which was replaced every other day. Cells were incubated at 37°C, 5 % CO_2_ and were passaged at 70-80 % confluency with 0.5 mM EDTA.

### PiggyBac-mediated NGN2 insertion into iPSCs

We generated *NGN2*-expressing iPSCs via piggyBac-mediated integration of a BFP-containing, doxycycline-inducible *NGN2*. iPSCs were washed with PBS and dissociated with Accutase (Gibco) and plated onto Geltrex-coated plates at the desired ratio (typically 1.5 × 10^6^ cells/well of a 6-well plate) in Stemflex plus 10 µM Y-2763 (Tocris). Once cells had adhered, cells were washed with PBS and transfected with constructs containing the donor doxycycline-inducible *NGN2* and a BFP fluorescent reporter flanked by transposon terminal repeats (Addgene: 172115) [7] and the piggyBac transposase. Cells were transfected with 2 µg total plasmid DNA (2:1 ratio of donor:tranposase) with Lipofectamine Stem, as per manufacturer’s instructions. 48 hours post transfection, media was changed to fresh media supplemented with 0.5 µg/mL puromycin for 3 days to select for those that had successfully integrated the donor plasmid. To ensure a pure population of NGN2-expressing cells, cells were subjected to fluorescence activated cell sorting (FACS) using BFP.

### iPSC differentiation in iLMNs

To generate lower-motor neuron like iPSC-neurons (iLMNs), transcription factor mediated differentiation was coupled with small molecule patterning. Detailed induction protocols for iLMNs can be found at Limone et al, 2023. Briefly, on day 0 cells were washed with PBS, lifted with Accutase (Gibco) and plated onto Geltrex-coated plates at the desired ratio (typically 400,000 cells/well of a 6-well plate) in Stemflex plus 10 µM Y-2763 (Tocris). On days 1-4, media was changed to N2 media consisting of DMEM-F12 (Gibco), 1X Glutamax (Gibco), 1X NEAA (Gibco), 1X N2 supplement (Gibco). On day 4, cells were lifted with Accutase and plated at the desired ratio (typically 800,000 cells/well of a 6-well plate) onto poly-L-ornithine (Merck) and laminin (Sigma)-coated plates in NB media, consisting of Neurobasal (Gibco), 1X Glutamax (Gibco), 1X NEAA (Gibco), N2 supplement (Gibco), B27 supplement (Gibco), 1X Culture One (Gibco) and 10 ng/mL neurotrophic factors (BDNF, GDNF, CNTF, PeproTech). Cells were maintained in NB media until day 8, and from then, media was replaced with fresh NB media without Culture One and 1/3 media changes were performed once a week. Cells were exposed to 10 µM SB431542 (S4317, Sigma) and 100 nM LDN-193189 (040019, Stemgent) on days 1-4, 1 µM retinoic acid (R2625, Merck) and 1 µM SAG (S7779-SEL, Stratech) on days 1-8, and 2 µg/mL doxycycline (D9891, Sigma) on days 2-8.

### Reverse-transcription quantitative PCR (RT-qPCR)

RNA was extracted from cell pellets using the ReliaPrep™ RNA cell kit (Promega) as per manufacturer’s instructions. For reverse transcription, 1 µg of RNA was mixed with SuperScript IV VILO Master Mix to produce cDNA as per the manufacturer’s instructions. Samples were incubated in a thermocycler with the following protocol: 25°C for 10 minutes, 50°C for 10 minutes, 85°C for 5 mins. Quantitative PCR (qPCR) was performed using SYBR Green Master Mix (Applied Biosystems) for all qPCRs except for *C9orf72* variant 2 where TaqMan probes were used. Primers were used at a final concentration of 1 µM with 5 ng cDNA per reaction in a 96-well plate. Signal was measured using the LightCycler (Roche) and samples were measured in triplicate and values were normalised to *GAPDH*, the housekeeping control. Transcripts were amplified with the following primers; *MAP2* (ggatatgcgctgattcttca, ctttccgttcatctgccatt), *HB9* (gtccaccgcgggcatgatcc, tcttcacctgggtctcggtgagc; Shimojo et al., 2015), *OCT4* (atgcattcaaactgaggtgcctgc, aacttcaccttccctcgaacgagt), Repeat-containing/Intron 1a (ccccactacttgctctcaca, cggttgtttccctccttgtt (Kempthorne et al 2024); Variant 2 (gcggtggcgagtggatat, tgggcaaagagtcgacatca; /56-FAM/atttggata/ZEN/atgtgacagttgg Fratta et al., 2013), Total *C9orf72* (actggaatggggatcgcagca, accctgatcttccattctctctgtgcc; Liu et al., 2014), *GAPDH* (gagtcaacggatttggtcat, ttgattttggagggatctcg).

### Repeat-primed PCR (rpPCR)

A repeat-primed PCR (rpPCR) reaction was performed on genomic DNA in the presence of 0.9 mM betaine, 7 % dimethyl sulfoxide, 0.9 mM MgCl2, 0.18 mM 7-deaza-dGTP and 1X Roche FastStart Master Mix, and is described previously (Renton et al., 2011). PCR products were mixed with HiDi Formamide (ABI: 4311320) and Liz500 standard (ABI: 4322682) and were analysed on an ABI3730 DNA Analyzer and visualized using GeneMapper software.

### Southern blotting

Southern blot analysis was performed on genomic DNA and is described previously [1]. Briefly, genomic DNA was extracted from iPSCs using the DNeasy kit (Qiagen) as per manufacturer’s instructions and digested overnight at 37°C with AluI/DdeI. 12 µg DNA was electrophoresed on a 0.8 % agarose gel alongside the DIG-labelled DNA molecular weight marker VII (11669940910, Sigma). The DNA was transferred overnight to a positively charged nylon membrane (1141724001, Sigma) by capillary blotting and was crossed-linked the following morning by UV. The membrane was prehybridized in Roche DIG Easy Hyb solution (11796895001, Sigma) with the addition of 100 µg/ml denatured salmon sperm (15632011, ThermoFisher) for 3 hours at 48°C. The membrane was probed overnight at 48°C with 10 ng/ml of a DIG labelled oligonucleotide comprising of five hexanucleotide repeats (GGGGCC)5 (Integrated DNA technologies). Following hybridization, the membrane was first washed in a 2 X saline-sodium citrate (SSC) and 0.5 % sodium dodecyl sulphate for 15 minutes while the hybridization oven ramped from 48°C to 65°C. The membrane was then further washed in 2 X SSC and 0.5 % SDS for 15 minutes, followed by 15 minute washes in 0.5 X SSC and 0.5 % SDS and 0.15 X SSC and 0.5 % SDS. Antibody detection was conducted following the DIG Application Manual protocol (Roche Applied Science) using the DIG wash and block buffer set (11585762001, Roche) and ready-to-use CSPD (11755633001, Sigma). Signals were visualised using ChemiDoc XRS+ system (BioRad). Membranes were exposed for 45 minutes with images collected every 4-5 minutes. Hexanucleotide repeat expansions were sized using Image Lab 6.1 software (BioRad) by comparison to DIG-labelled DNA molecular weight marker VII and subtraction of the genomic non repeat region of C9 flanking the AluI/DdeI restriction cut sites (156bp).

### Immunoblotting

To extract protein, cell pellets were lysed with lysis buffer (2 % SDS + protease inhibitors in RIPA buffer) for 10 minutes on ice before sonication for 3 × 5 seconds at 30 amp at 4°C. Samples were then centrifuged at 17,000 xg at 4°C for 20 minutes and the supernatant was collected and stored at −80°C. Protein concentration was determined using a BCA protein assay kit (ThermoFisher) as per manufacturer’s instructions. Protein extracts were denatured at 99°C for 10 minutes in 1 X Laemmli buffer (Alfa Aesar) and separated on a NuPAGE™ 4 – 12 % bis-tris gel at 100 V in NuPage MES buffer (ThermoFisher) for 1.5 hours. Gels were transferred to nitrocellulose membrane (Bio-Rad Laboratories) and membranes were blocked in 5 % milk in PBS-T (0.1 % Tween-20) for 1 hour and incubated overnight at 4°C with the following primary antibodies diluted in 5 % milk PBS-T: C9orf72 (GTX634482, GeneTex, 1:1000) and Calnexin (sc-6465, Santa Cruz Biotechnology, 1:1000). Membranes were then washed 3 times in PBS-T prior to 1 hour incubation with secondary HRP-conjugated antibodies diluted in 5 % milk PBS-T at room temperature. After 3 washes in PBS-T, membranes were treated with Immobilon Forte Western HRP substrate (Millipore) and imaged using the Amersham imager 680, GE life sciences) and quantifications were performed using ImageJ software.

### MSD immunoassays

The Meso Scale Discovery (MSD) immunoassay was performed to detected poly(GA) or poly(GP) expression levels and is described previously [25,26]. Briefly, samples were prepared to a unified concentration of 0.5 mg/mL in lysis buffer (RIPA + 2 % SDS and protease inhibitors) and 25 µL were loaded in duplicate in the 96-well MSD immunoassay plate. The following antibodies were used: anti-poly(GA) (MABN889, Sigma) and anti-poly(GP) (GP658, custom-made from Eurogentec) and biotinylated, streptavidin-conjugated, sulfo-tagged anti-poly(GA) (GA 5F2*, a kind gift from Dieter Edbauer) and anti-poly(GP) (GP658*, custom-made from Eurogentec) as detector antibodies. Plates were read with MSD reading buffer (R92TC, MSD) using the MSD Sector Imager 2400.

### Immunostaining

iLMNs were plated in a 96-well PhenoPlate (PerkinElmer) and fixed on day 10. Briefly, on day of fixation, cells were washed with PBS and fixed using 4 % paraformaldehyde in PBS for 10 minutes at room temperature. After incubation, cells were blocked and permeablised in 10 % FBS and 0.25 % Triton-X-100 in PBS for 45 minutes at room temperature. Cells were washed 3 times in PBS and incubated with primary antibodies in 10 % FBS in PBS overnight at 4°C. The following primary antibodies were used: MAP2 (NB300-213, Novus, 1:2000), SMI-32 (N2912, Sigma, 1:100). After 3 PBS washes, cells were incubated with secondary antibodies in 10 % FBS in PBS. Cells were washed 3 times in PBS prior to incubation with Hoescht (1:5000) for 10 minutes at room temperature. Cells were imaged using the automated Opera Phenix TM high-throughput confocal imaging platform (PerkinElmer). The Harmony high-content analysis software (Harmony v4.8, PerkinElmer) was used to quantify the percentage of SMI-32 expressing cells.

### RNA foci detection

Cells were fixed as described above and then dehydrated via 70 % ethanol followed by 100 % ethanol washes and frozen at −80°C in 100 % ethanol until needed. On the day of the experiment, cells were rehydrated in 70 % ethanol before being permeabilised by incubation with 0.2 % Triton-X-100 in PBS for 10 minutes at room temperature. Samples were then blocked in hybridisation solution (40 % formamide, 2 X SSC, 10 % dextran sulphate, 2 mM vanadyl ribonucleoside complex) at 66°C for 45 minutes. To visualise foci, locked nucleic acid (LNA) probes (Qiagen) targeting sense RNA foci (5’ TYE563-labelled CCCCGGCCCCGGCCCC) were used at 40 nM in hybridisation solution. Cells were incubated in probe-hybridisation solution in the dark at 66°C for 3 hours. Following incubation, cells were washed with 0.2 % Triton-X100 in 2 X saline-sodium citrate (SSC) for 5 minutes at room temperature followed by incubation at 66°C for 30 minutes. Cells were then washed in 0.2 X SSC at 66°C for an additional 30 minutes prior to incubation overnight with an anti-DIG antibody conjugated to Alexa-647 (in 0.2 X SSC with 1 % BSA, 1:1000). A final 20-minute wash with 0.2 X SSC was performed at room temperature before incubation with Hoechst (1:5000) for 10 minutes. Hoechst solution was removed and cells were left in 0.2 X SSC at 4°C before imaging on the Opera Phenix High Content Imager.

## Acknowledgements and funding

This work is supported by the UK Dementia Research Institute (award number UK DRI-1006) through UK DRI Ltd, principally funded by the Medical Research Council and an Eisai funded Leonard Wolfson Experimental Neurology Centre PhD Studentship to R.C. Engineering of repeat constructs was funded by the Medical Research Council (award number MC_EX_MR/N501931/1). The generation of C9 repeat expansion clones was supported by NIA contracts to the Jackson Laboratory for Genomic Medicine for the iPSC Neurodegenerative Disease Initiative (iNDI) project. We gratefully acknowledge the contribution of the Scientific Services at the Jackson Laboratory for work described in this publication, including the Cellular Engineering service for SNP array analysis of iPS cell clones and the Genome Technologies service for expert assistance with Cas9 targeted nanopore sequencing. We thank Quintara for Sanger sequencing services and Cell Line Genetics (Madison, WI) for G-band karyotyping.

## Supplementary Figures

**Supplemental Figure 1.**
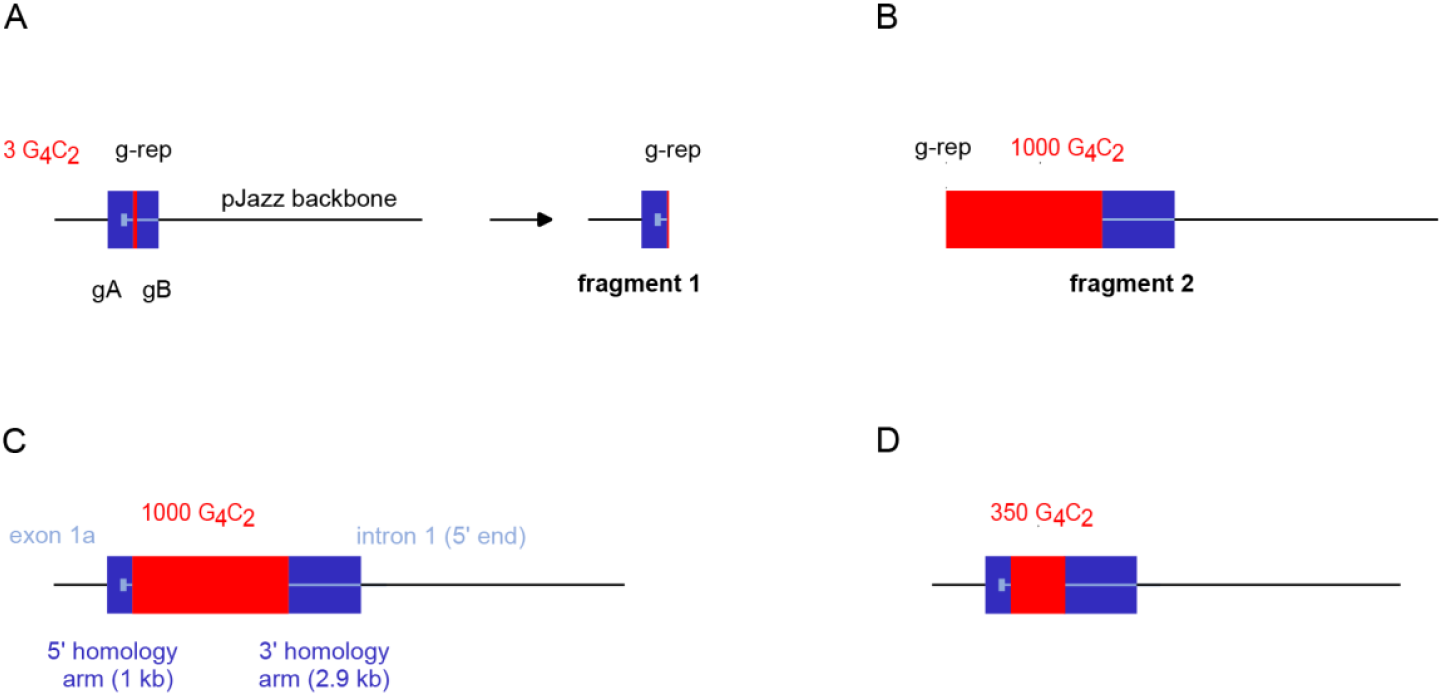
Assembly of the human iPSC targeting construct. (**A**) A ~2kb fragment of human *C9orf72* including the G_4_C_2_ region with 3 repeats was first isolated via CRISPR-Cas9 digestion from a human C9orf72 locus BAC clone (RP11-27J8), using sgRNA guides A and B (gA and gB), and blunt end cloned into the pJazz-OK backbone. CRISPR-Cas9 digestion with an sgRNA guide targeting within the 5’ end of the repeat sequence (g-rep) was then used to generate fragment 1, constituting the 5’ homology arm (~1kb) together with the short arm of the pJazz vector. (**B**) Fragment 2 was derived from an existing construct originally designed for generating mouse models (from ‘pJazz-V4’ [10]), and harbours 1000 G_4_C_2_ repeats plus 2.9 kb of flanking human sequence (3’ homology arm), together with the long arm of the pJazz vector; and was isolated from its parent vector via CRISPR-Cas9 digestion with sgRNA guide g-rep, cutting within the 5’ end of the long repeat. (**C**) Fragments 1 and 2 were ligated together to form a targeting construct with a seamless 1000 G_4_C_2_ repeat expansion within the endogenous human genomic sequence. (**D**) A natural retraction of the repeat during the final cloning step additionally yielded a 350-repeat targeting construct. The 350-repeat targeting construct was taken forward, as it was non-toxic, in contrast to the 1000-repeat construct, which resulted in cell death following nucleofection. sgRNA gA: AACGTTTTAATCATTCACCG; sgRNA gB: TTTCTGAATACAAAGCCTGG; sgRNA g-rep: AGGAGTCGCGCGCTAGGGGC.

**Supplemental Figure 2.**
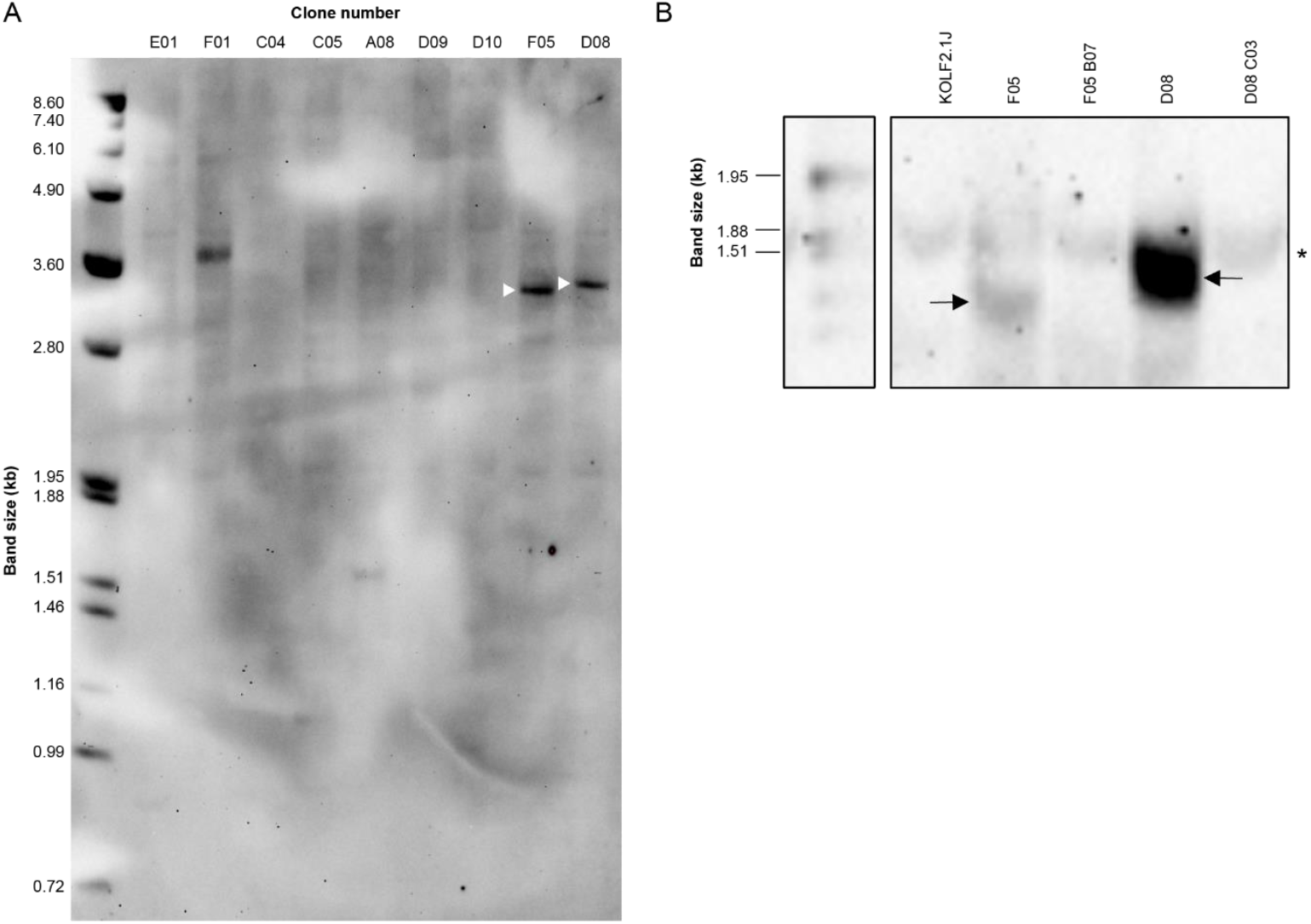
Southern blotting of repeat knock-in and revertant lines. (**A**) Southern blot of GGGGCC repeats in initial iPSC knock-in clones, demonstrating successful integration of expanded repeats in clones F05 and D08 (white arrows). (**B**) Southern blot of F05 and D08, with their respective revertant lines and the parental KOLF2.1J shows repeat expansions of approximately 200 repeats in F05 and D08 and successful removal in the revertant lines, *non-specific band.

**Supplemental Figure 3.**
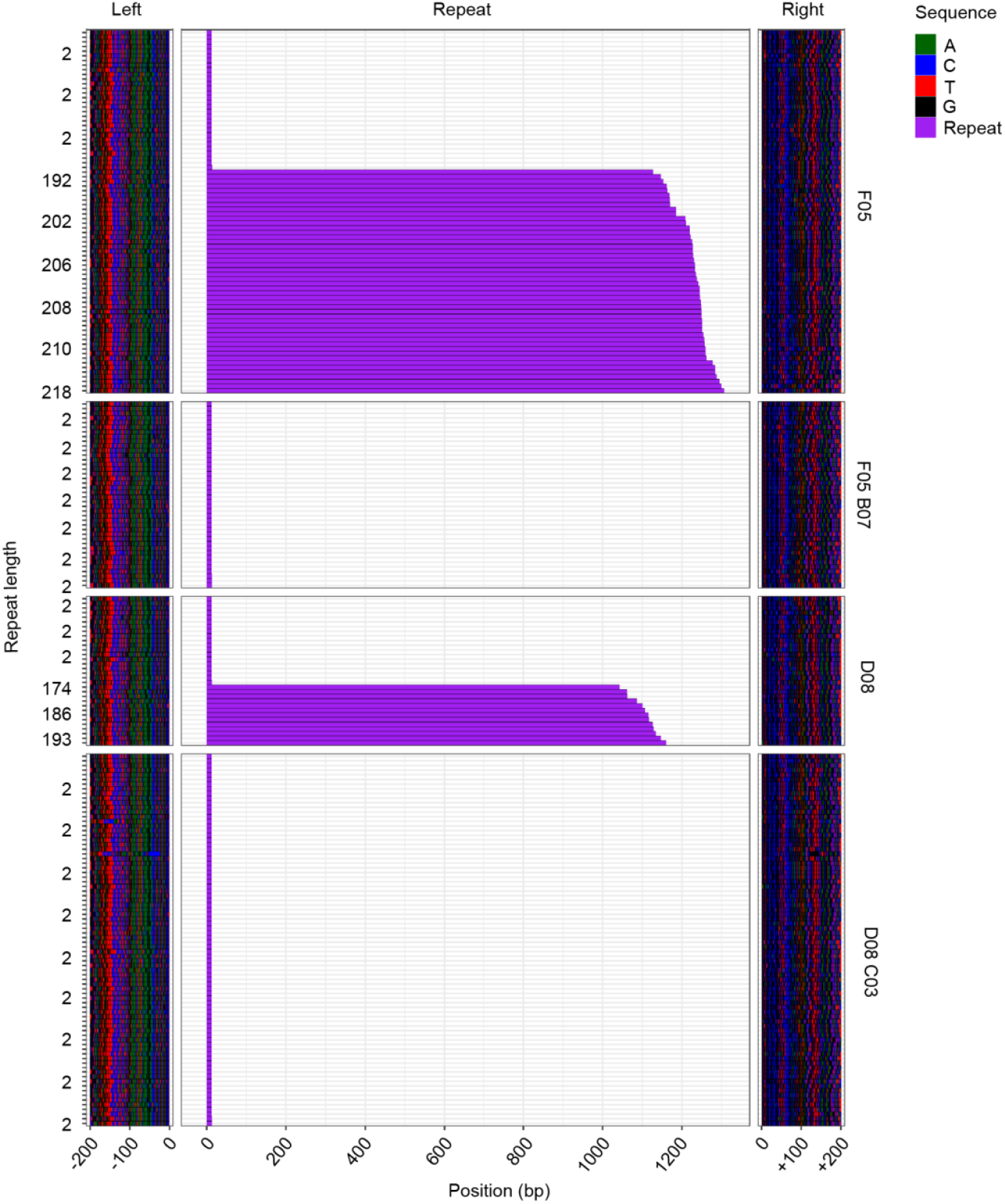
Waterfall plots showing repeat length distribution and flanking sequence for each *C9orf72* repeat knock-in and revertant line. ONT long-read *C9orf72* sequences were trimmed to 200 bases flanking the hexanucleotide repeat. Each row represents a read, with the x-axis indicating base position along the read and the y-axis giving hexanucleotide repeat length. The plot is divided into three horizontal facets; *Left, Repeat* and *Right*, corresponding to the 200-base flanking regions and the repeat region. Coloured tiles represent individual nucleotide bases, with the colour scheme detailed in the key.

